# Optimization of whole-genome sequencing of *Plasmodium falciparum* from low-density dried blood spot samples

**DOI:** 10.1101/835389

**Authors:** Noam B. Teyssier, Anna Chen, Elias M Duarte, Rene Sit, Bryan Greenhouse, Sofonias K. Tessema

## Abstract

**Background:** Whole-genome sequencing (WGS) is becoming increasingly useful to study the biology, epidemiology, and ecology of malaria parasites. Despite ease of sampling, DNA extracted from dried blood spots (DBS) has a high ratio of human DNA compared to parasite DNA, which poses a challenge for downstream genetic analyses. We evaluated the effects of multiple methods for DNA extraction, digestion of methylated DNA, and amplification on the quality and fidelity of WGS data recovered from DBS.

**Results:** At 100 parasites/μL, Chelex-Tween-McrBC samples had higher coverage (5X depth = 93% genome) than QIAamp extracted samples (5X depth = 76% genome). The two evaluated sWGA primer sets showed minor differences in overall genome coverage and SNP concordance, with a newly proposed combination of 20 primers showing a modest improvement in coverage over those previously published.

**Conclusions:** Overall, Tween-Chelex extracted samples that were treated with McrBC digestion and are amplified using 6A10AD sWGA conditions had minimal dropout rate, higher percentages of coverage at higher depth, and more accurate SNP concordance than QiaAMP extracted samples. These findings extend the results of previously reported methods, making whole genome sequencing accessible to a larger number of low density samples that are commonly encountered in cross-sectional surveys.

## Background

Whole genome sequence (WGS) data provide a complete picture of a pathogen genome and are transforming molecular epidemiology [1]. In the past few years, the availability of WGS data from field collected malaria samples has grown along with the development of more sensitive, faster and lower-cost sequencing technologies. Until recently, generating these sequence data required collection of large volumes of venous blood, leukocyte depletion at the time of sample collection, and storage in the field - tasks often challenging to perform in resource-limited settings [2,3]. In contrast, collection of dried blood spots (DBS) requires small sample volumes and such samples are easily stored and transported. However, the low volumes of blood present in DBS often result in low quality and quantity of parasite DNA, particularly relative to the overwhelming proportion of human DNA in the sample. In order to address these challenges, several studies have utilized different enrichment methods such as non-selective whole genome amplification (WGA) [4,5]; hybrid selection [6–9]; enzymatic digestion of host DNA using the MspJI family of restriction enzymes [10,11] and selective whole genome amplification (sWGA) [11–14]. Recently, collection of leukocyte-depleted dried erythrocyte spots has also showed promise to provide parasite enriched samples for WGS [15].

Unlike other enrichment methods, sWGA requires relatively small amounts of starting material and provides relatively high quality data for a low-cost - enabling WGS of field samples that would otherwise contain insufficient genetic material for *P. falciparum* [13], *P. vivax* [11] *and P. Knowlesi* [14]. However, it is possible that current sWGA protocols can be improved to increase coverage, sensitivity, and reproducibility. For example, two recent studies that performed sWGA on clinical samples using two different sets of sWGA primers, reported an average 55% of reads mapped to the *P. falciparum* genome with 48% of the genome covered at ≥5X in 24 samples with an average parasite density of 73,601 parasites per µL [13,16]. However, Oyola *et al.*, established a parasitaemia threshold of 0.03% (>1200 parasites/µL) to obtain at least 50% of the genome at 5X in clinical samples. Similar level of coverage was reported in a *P. vivax* specific sWGA [11], highlighting the need to improve these protocols for DBS samples.

The efficiency of sWGA could be potentially be improved by standardized comparison and optimization of different DNA extraction methods from DBS; identifying optimal sWGA primer sets that provide higher yield with reduced bias; and performing enzymatic cleavage of human DNA prior to sWGA [10]. In this study, we compared four DNA extraction methods followed by either enzymatic cleavage of human DNA or no cleavage, and amplification with WGA or sWGA using different primer sets. The combination of conditions were compared in standardized samples across a range of low parasite densities extracted from DBS, and the resulting WGS data were compared in terms of coverage of the parasite genome, dropout rate, and the concordance of called variants.

## Results

### Experimental design

Dried blood spots (DBS) with densities from 10 to 10,000 parasites/μL blood were extracted using four different DNA extraction methods and then amplified by two different sets of sWGA primers or random hexamer primers with or without a prior treatment of input DNA with McrBC, an endonuclease which cleaves human DNA containing methyl cytosine [17] (Figure 1). The recovery and quality of WGS were compared between the different combinations of methods.

**Figure 1.**
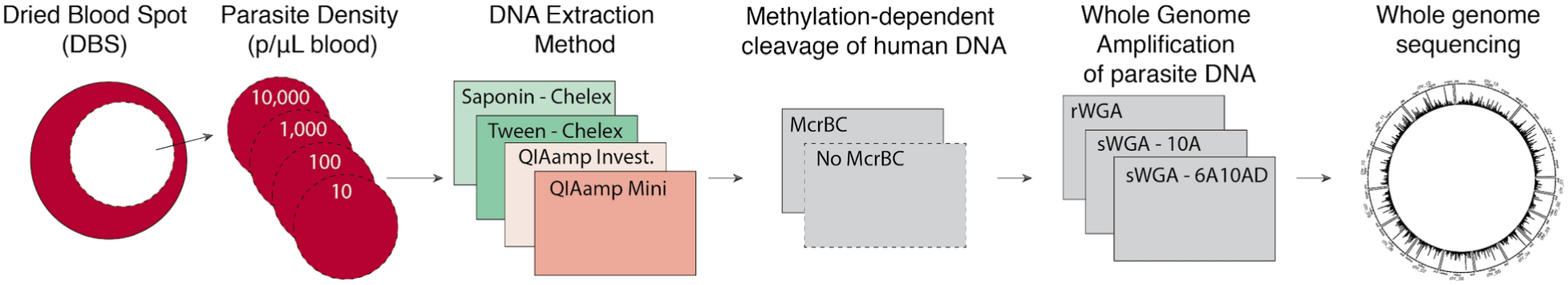
Overview of the experimental workflow. Various combinations of DNA extraction, methylation dependent cleavage of human DNA, and whole genome amplification were compared. Abbreviations: rWGA is whole genome amplification using random hexamers. sWGA is selective whole genome amplification using one of two sets of P. falciparum specific primers (10A and 6A10AD).

### Tween-Chelex extraction yielded the highest coverage in low density samples

The recovery and coverage of WGS for DNA extraction methods across parasite densities were investigated. Four methods were evaluated, two of which were spin column based and two Chelex based. DBS extracted with the Saponin-Chelex method had the lowest performance, with only 13% of reads mapping to the *P. falciparum* core genome and poor genome coverage (1X coverage ranging from 2% to 50%) across all parasite densities. This extraction method was not investigated further. QiaAMP Mini and QiaAMP Investigator spin column kits performed similarly in terms of the percentage of reads that mapped to *P. falciparum* regardless of parasitemia. However, the QiaAMP Investigator kit had a 5% higher dropout rate than QiaAMP Mini at 1000 parasites/μL (15% vs 20%) and 20% higher at 100 parasites/μL (21% vs 42%), and was also not investigated further. The performance of Tween-Chelex extraction and QiaAMP Mini kits were evaluated in further detail.

Extraction with QiaAMP Mini resulted in a consistently higher proportion of reads mapping to *P. falciparum* (Figure 2A). However, the higher mapping rate did not translate into better sequence coverage across the *P. falciparum* core genome (Figure 2B). Samples extracted using the Tween-Chelex method consistently resulted in a greater proportion of the genome covered at 5X depth than samples extracted via spin columns (e.g., 89% vs 72% at 100 p/μL). Similarly, DBS extracted via spin columns had higher dropout rates than those extracted with Tween-Chelex (e.g., 12% vs. 2% respectively at 100 parasites/μL) (Figure 3). In order to evaluate the accuracy of the mapped reads, the overall SNP concordance was compared for known SNP positions and *de novo* SNP calls to the reference 3D7 genome. Both extraction methods achieved high concordance, with Tween-Chelex outperforming QiaAMP for lower density samples (Figure 4). Overall, the Tween-Chelex extraction method had a lower dropout rate, higher genome-wide coverage and higher rate of SNP concordance than the QiaAMP mini kit on low density DBS samples for a given number of total reads, despite having a lower fraction of these reads mapping to the *P. falciparum* core genome.

**Figure 2.**
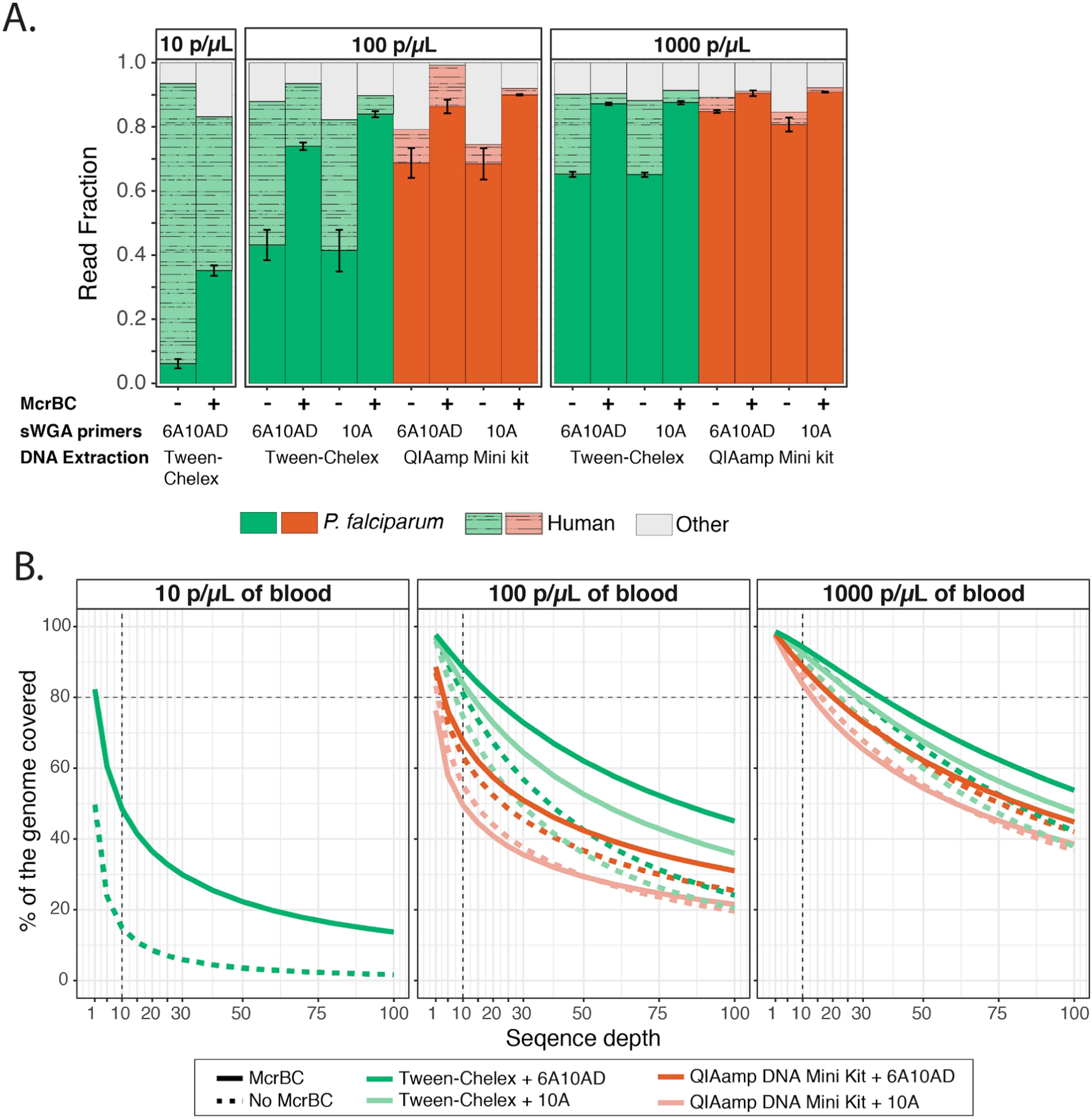
Comparisons of read mapping and genome coverage. **A.** Proportion of reads that mapped to to either the reference P. falciparum 3D7 or hg19 human genome. Unmapped reads are indicated as other. Error bars indicate standard error between 2 experimental replicates. **B.** The percentage of the core P. falciparum genome covered by a minimum depth is shown.

**Figure 3.**
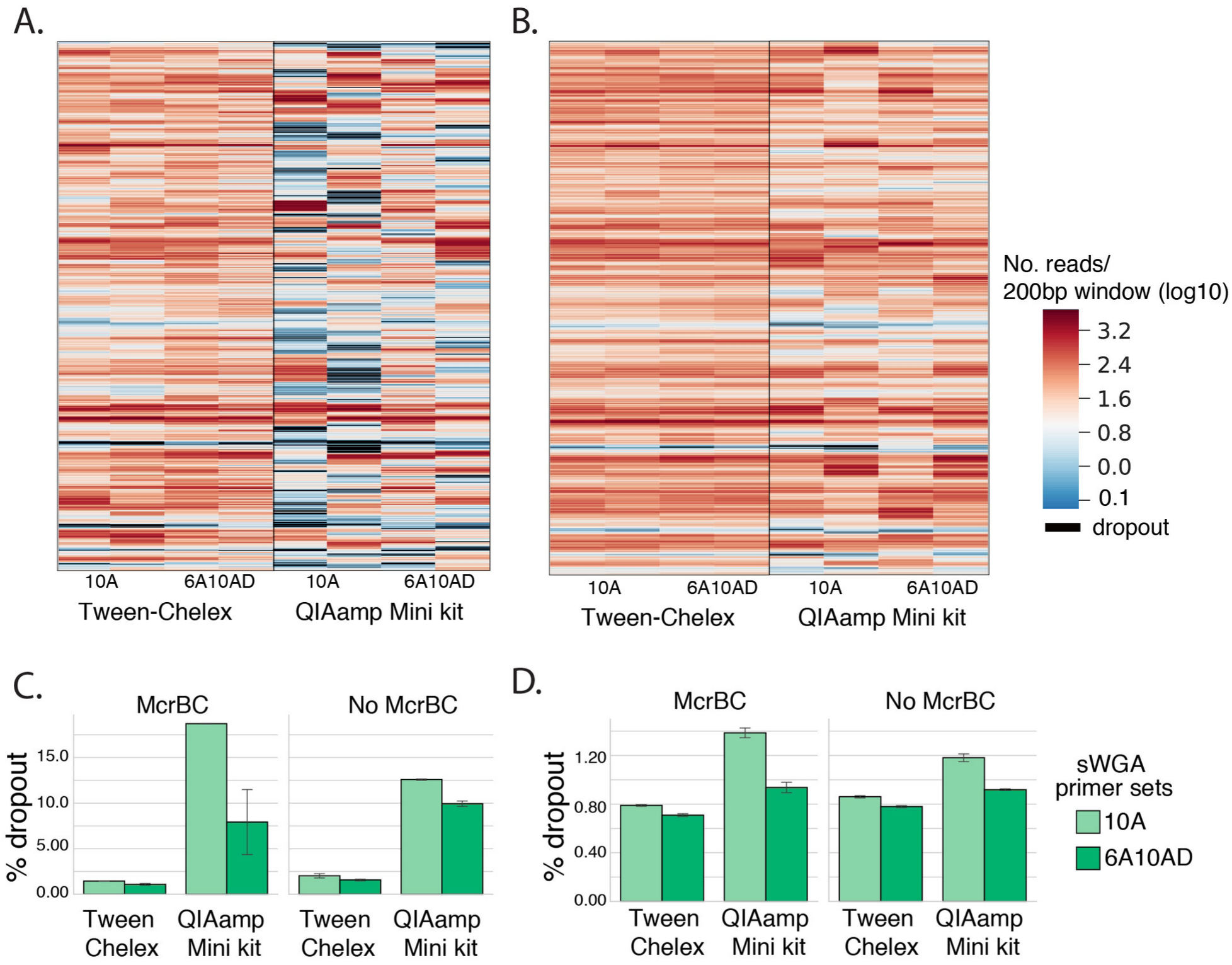
Comparisons of genome coverage and dropout rate. Representative coverage depth of chromosome 7 for McrBC followed by sWGA digested samples at 100 (A) and 1000 (B) parasites/μL. Coverage was determined as the mean number of reads in 200bp interval windows of the core genome. Panels C and D show dropout rate, i.e., the percentage of the genome that was not covered across all chromosomes in 100 (C) and 1000 (D) parasites/μL.

**Figure 4.**
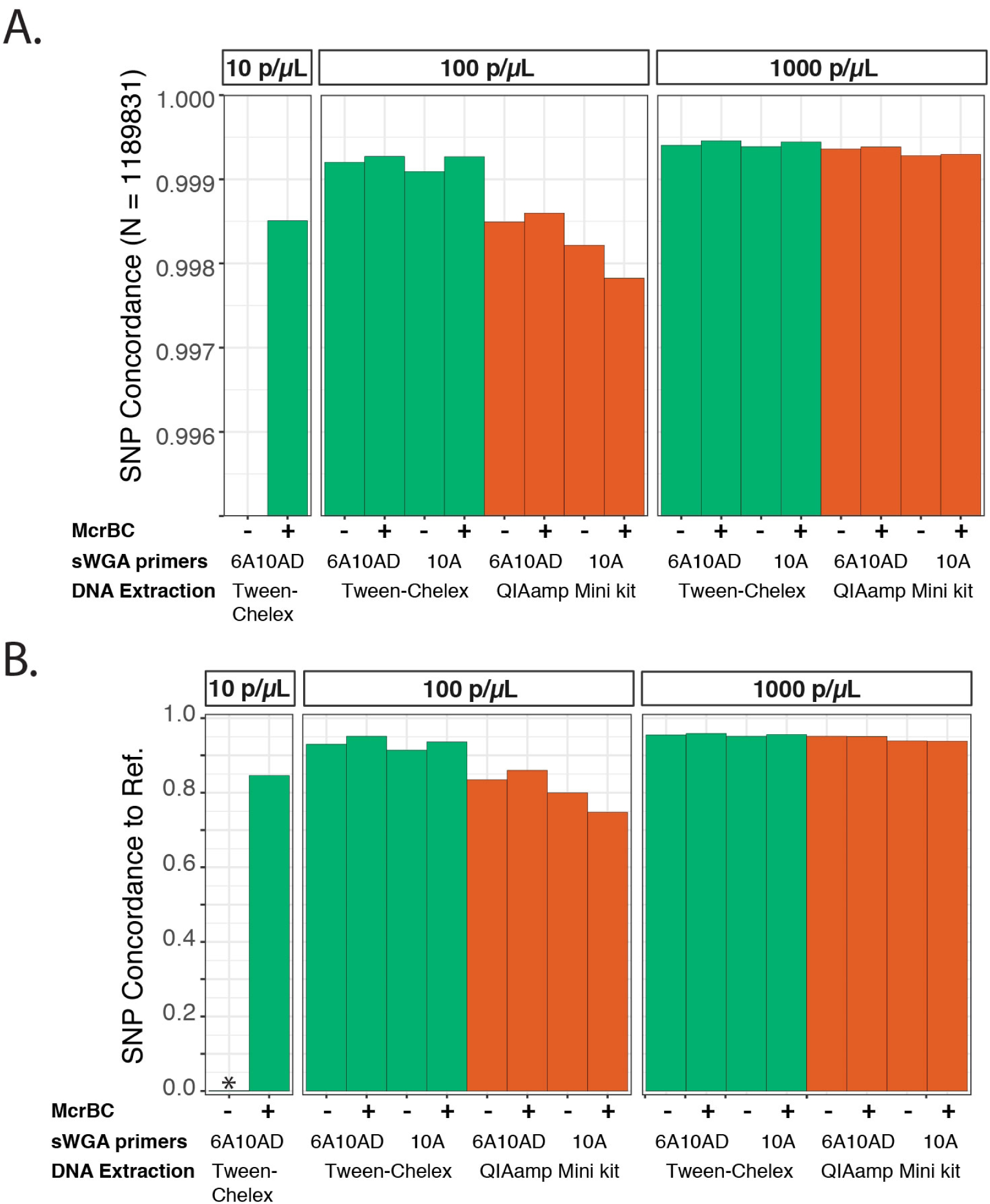
Comparison of variant call concordance between experimental conditions and shotgun sequenced reference strains. **A.** High quality biallelic SNP positions from pf3k_5.1 (https://www.malariagen.net/projects/Pf3k) were compared, as in Oyola *et al.*, 2016 [13]. **B.** High quality homozygous non-reference variant sites called de novo were compared. SNPs were not called for 10 parasites/μL samples without the addition of McrBC due to the lack of sufficient read depth.

### McrBC improves sensitivity and genome coverage in low density samples

Whole genome amplification has been shown to be a successful strategy for increasing the amount of material available for WGS of *P. falciparum* from DBS samples. This study compared performance for two sets of *P. falciparum* specific sWGA primer sets, 10A and 6A10AD, and a random hexamer-based WGA. Overall, the two sWGA primer sets performed similarly with respect to the percentage of reads that mapped to *P. falciparum* (Figure 2B) and genotype concordance (Figure 3). However, the 6A10AD primer set resulted in moderately higher breadth and depth of coverage across extraction methods and parasite densities (Figure 2C).

To test an alternative enrichment strategy to sWGA, WGA with random hexamers preceded by McrBC treatment was performed, resulting in a significantly lower genome coverage than sWGA. However, performing sWGA on McrBC treated samples yielded a substantial increase in the depth of genome coverage that is most marked at the lowest parasite densities extracted with Tween-Chelex. After McrBC treatment, the proportion of reads that mapped to the human genome dropped considerably at 100 parasites/μL (Tween-Chelex: 43% to 5%), and 10 parasites/μL (Tween-Chelex: 87% to 47%), dramatically improving usability and cost efficiency of the assay (Figure 2A). Conversely, McrBC improved *P. falciparum* genome coverage, with a more pronounced increase in Tween-Chelex than QiaAMP mini extracted samples. Samples at 10 parasites/μL had a substantial increase in the percentage of the genome covered at 5X from 23% to 60% (Figure 2B).

## Discussion

The ability to recover high quality sequencing data from low-density dried blood spots has useful and immediate implications for public health - allowing sample collection to be performed in a cost effective and scalable manner. In this study, a comprehensive evaluation was performed on the effects of DNA extraction method, McrBC based digestion of human DNA, and sWGA conditions across a range of parasite densities. Tween-Chelex extraction, digestion with McrBC, and sWGA using a new combination of 20 previously published primers [12,13] and an AT-biased dNTP mixture provided the highest yield and most reliable coverage, allowing whole genome sequencing data to be obtained from dried blood spots containing as few as 10 parasites per µL of blood. These findings extend the results of previously reported methods, making whole genome sequencing accessible to a larger number of low density samples that are common in cross-sectional surveys.

The data show that recovery of WGS from DBS samples with 10 parasites/µL of blood is possible, but the quality and accuracy was higher for samples containing at least 100 parasites/µL (e.g. 93% of the genome at 5X coverage). McrBC treatment prior to sWGA allows for even greater recovery across all methods evaluated, and improved performance in all metrics tested. Interestingly, QiaAMP extraction resulted in a higher proportion of total reads mapping to the *P. falciparum* genome than Tween-Chelex extraction, but nonetheless resulted in lower coverage, higher drop-out, and lower SNP concordance. This apparent discrepancy could stem from preferential recovery of specific regions of the genome by spin columns but more even recovery by Tween-Chelex, and can be seen as a compensating increase in a small part of the genome being covered at higher depth.

The factor evaluated with the least effect on sequence quality and recovery was the sWGA primer set, but there were still modest benefits to using a more expanded primer set with AT-biased dNTPs (6A10AD). These effects were present independent of extraction and digestion method, and showed consistent improvements over the previously published 10A primer set.

## Conclusions

This study showed that Tween-Chelex extracted samples treated with McrBC digestion and amplified using 6A10AD sWGA conditions had higher performance with respect to minimal genome dropout, higher percentages of coverage at higher depth, and more accurate SNP concordance with respect to reference strains. Tween-Chelex extraction has the added benefit of being cheaper than spin columns, allowing researchers and elimination programs to scale analyses with respect to dollar per base coverage. McrBC digestion prior to sWGA provided a significant improvement over non-digested samples extracted with Tween-Chelex, most dramatic in the lowest parasite density samples. There are numerous logistical advantages of DBS collection over venous blood, including a lower barrier of training for sample collection, less blood to collect per sample, and no specialized transportation requirements, allowing researchers and control programs to incorporate whole genome sequencing, including samples as low as 10 parasites per µL, into research and surveillance activities where such data will be useful.

## Methods

### Mock DBS samples

Mock DBS samples were made by mixing synchronized, ring stage cultured *P. falciparum* parasites with uninfected human whole blood to obtain a range of parasite densities (10, 100, 1000 and 10000 parasites per μL of blood). DBS samples were stored at -20°C until processing.

### DNA extraction and quantitative PCR

DNA was extracted from a single 6mm hole-punch using four different extraction methods. i) Saponin-Chelex as described previously [18]; ii) a modified Tween-Chelex described below; iii) QIAamp DNA Mini Kit (Qiagen, California, United States) following the manufacturer’s recommendations and iv) QIAamp DNA Investigator Kit (Qiagen, California, United States) as described elsewhere [13].

Tween-Chelex extraction was conducted by modifying the Saponin-Chelex extraction method. DBS were punched using a 6mm hole-puncher in to 1.5ml microcentrifuge tubes. 1mL of 0.5% Tween 20 (catalogue # P1379, Sigma Aldrich) in 1X PBS was added into the tube containing DBS punches and incubated overnight at 4°C. The samples were briefly centrifuged, Tween-PBS was removed and the punches were washed with 1mL of 1X PBS and incubated for 30 minutes at 4°C. The samples were briefly centrifuged, PBS removed and 150μL of 10% Chelex 100 resin (catalogue # 1422822, Bio-Rad Laboratories) in water were added to each sample, ensuring the DBS punches were covered with the Chelex solution and incubated for 10 minutes at 95°C. The tubes were centrifuged at 15,000rpm for 10 minutes and the supernatant was transferred to 0.6mL microcentrifuge tubes and centrifuged at 15,000rpm for 5 minutes. The extracted DNA was then transferred to a 96-well plate and stored at -20°C until processing. The parasite densities were confirmed using *var*-ATS ultra-sensitive qPCR as described previously [19].

### McrBC digestion of human DNA

For a subset of samples, extracted DNA was digested with McrBC (catalogue # M0272S, New England Biolabs) in a 30 μl reaction containing 20μL of extracted DNA, 10 units of McrBC, 1X NEBuffer 2, 0.5μl 100X BSA and 0.5μl 100X GTP. The samples were incubated at 37°C for 2 hrs followed by inactivation at 65°C for 20 minutes. The digested products were used as templates for whole-genome amplification.

### Selective Whole genome amplification (sWGA)

The sWGA reactions were performed using two sets of primers: 10A [13] and 6A10AD, a combination of 6A [12] and 10A [13] with an adjusted ratio of dNTPs. The sWGA reaction using the 6A10AD primer set was performed as described previously [13] except for the adjusted proportion of nucleotides in the dNTP mix (i.e. 70% AT and 30% GC), similar to the composition of the malaria genome. The master mix and reaction condition for the modified sWGA protocol is shown in Table S1. A template DNA volume of 20uL with an estimated 16-16000 parasite genomes (Table S2) per sWGA reaction was used. For the non-selective whole genome amplification, random hexamer primers were used following the manufacturer’s instructions for the GenomiPhi V2 DNA Amplification Kit (catalogue # 45-001-221, GE Healthcare Life Sciences).

### Whole-genome sequencing

The whole genome amplification product was purified with SPRI magnetic beads (catalogue #65152105050250) to remove unbound primers, primer dimers, and other impurities. Sequencing libraries were prepared using the NEBNext® Ultra™ II DNA Library Prep Kit (catalogue #E7103) following manufacturer’s instructions. Samples were barcoded, purified, pooled and sequenced on the Illumina NextSeq550 or NovaSeq 6000 System using 150bp paired-end sequencing chemistry.

### Sequence analyses

Reads were demultiplexed, filtered by base quality and poly-g tails were clipped. The reads were then aligned to the *P. falciparum* 3D7 reference genome (version 3) with BWA-MEM with secondary alignments marked [20]. Sample base quality was recalibrated using Genome Analysis Toolkit (GATK) BaseRecalibrator and GATK ApplyBQSR with SNP locations from the Pf3k project (Data Release 5) used as a prior. The number of reads per sample were then downsampled to minimum total reads for equivalent comparisons. Percentiles of genome coverage were calculated using GATK CollectWgsMetrics across the core genome. Dropouts were evaluated by taking mean coverage over 200bp sliding windows across the core genome. Percentage of reads mapping to the core *P. falciparum* genome was calculated with samtools flagstat [21].

Variant calling and variant-filtering was conducted on the genome sequences following the Genome Analysis Toolkit (GATK) Best Practices [22] with minor modifications. Variants were called by sample across the core genome using gatk4 HaplotypeCaller (-ERC GVCF) then genotyped across all samples using gatk4 CombineGVCFs and gatk4 GenotypeGVCFs. The variants were recalibrated using gatk4 VariantRecalibrator and ApplyVQSR using data from the Pf3k project (Data Release 5) as a training set (annotations: QD, MQ, MQRankSum, ReadPosRankSum, FS, SOR). Variants were then filtered for passing VQSLOD and for biallelic SNPs. SNP concordance was measured with bcftools gtcheck, and gatk GenotypeConcordance.

## References

1. Tang P, Croxen MA, Hasan MR, Hsiao WWL, Hoang LM. Infection control in the new age of genomic epidemiology. Am J Infect Control. 2017;45:170–9.

2. Auburn S, Campino S, Clark TG, Djimde AA, Zongo I, Pinches R, et al. An effective method to purify Plasmodium falciparum DNA directly from clinical blood samples for whole genome high-throughput sequencing. PLoS One. 2011;6:e22213.

3. Venkatesan M, Amaratunga C, Campino S, Auburn S, Koch O, Lim P, et al. Using CF11 cellulose columns to inexpensively and effectively remove human DNA from Plasmodium falciparum-infected whole blood samples. Malar J. 2012;11:41.

4. Wang Y, Nair S, Nosten F, Anderson TJC. Multiple displacement amplification for malaria parasite DNA. J Parasitol. 2009;95:253–5.

5. Oyola SO, Manske M, Campino S, Claessens A, Hamilton WL, Kekre M, et al. Optimized whole-genome amplification strategy for extremely AT-biased template. DNA Res. 2014;21:661–71.

6. Melnikov A, Galinsky K, Rogov P, Fennell T, Van Tyne D, Russ C, et al. Hybrid selection for sequencing pathogen genomes from clinical samples. Genome Biol. 2011;12:R73.

7. Smith M, Campino S, Gu Y, Clark TG, Otto TD, Maslen G, et al. An In-Solution Hybridisation Method for the Isolation of Pathogen DNA from Human DNA-rich Clinical Samples for Analysis by NGS. Open Genomics J [Internet]. 2012;5. Available from: http://dx.doi.org/10.2174/1875693X01205010018

8. Bright AT, Tewhey R, Abeles S, Chuquiyauri R, Llanos-Cuentas A, Ferreira MU, et al. Whole genome sequencing analysis of Plasmodium vivax using whole genome capture. BMC Genomics. 2012;13:262.

9. Marciniak S, Prowse TL, Herring DA, Klunk J, Kuch M, Duggan AT, et al. Plasmodium falciparum malaria in 1st-2nd century CE southern Italy. Curr Biol. 2016;26:R1220–2.

10. Oyola SO, Gu Y, Manske M, Otto TD, O’Brien J, Alcock D, et al. Efficient depletion of host DNA contamination in malaria clinical sequencing. J Clin Microbiol. 2013;51:745–51.

11. Cowell AN, Loy DE, Sundararaman SA, Valdivia H, Fisch K, Lescano AG, et al. Selective Whole-Genome Amplification Is a Robust Method That Enables Scalable Whole-Genome Sequencing of Plasmodium vivax from Unprocessed Clinical Samples. MBio [Internet]. 2017;8. Available from: http://dx.doi.org/10.1128/mBio.02257-16

12. Sundararaman SA, Plenderleith LJ, Liu W, Loy DE, Learn GH, Li Y, et al. Genomes of cryptic chimpanzee Plasmodium species reveal key evolutionary events leading to human malaria. Nat Commun. 2016;7:11078.

13. Oyola SO, Ariani CV, Hamilton WL, Kekre M, Amenga-Etego LN, Ghansah A, et al. Whole genome sequencing of Plasmodium falciparum from dried blood spots using selective whole genome amplification. Malar J. 2016;15:597.

14. Benavente ED, Gomes AR, De Silva JR, Grigg M, Walker H, Barber BE, et al. Whole genome sequencing of amplified Plasmodium knowlesi DNA from unprocessed blood reveals genetic exchange events between Malaysian Peninsular and Borneo subpopulations. Sci Rep. 2019;9:9873.

15. Nag S, Kofoed P-E, Ursing J, Lemvigh CK, Allesøe RL, Rodrigues A, et al. Direct whole-genome sequencing of Plasmodium falciparum specimens from dried erythrocyte spots. Malar J. 2018;17:91.

16. Guggisberg AM, Sundararaman SA, Lanaspa M, Moraleda C, González R, Mayor A, et al. Whole-Genome Sequencing to Evaluate the Resistance Landscape Following Antimalarial Treatment Failure With Fosmidomycin-Clindamycin. J Infect Dis. 2016;214:1085–91.

17. Panne D, Raleigh EA, Bickle TA. The McrBC endonuclease translocates DNA in a reaction dependent on GTP hydrolysis. J Mol Biol. 1999;290:49–60.

18. Plowe CV, Djimde A, Bouare M, Doumbo O, Wellems TE. Pyrimethamine and proguanil resistance-conferring mutations in Plasmodium falciparum dihydrofolate reductase: polymerase chain reaction methods for surveillance in Africa. Am J Trop Med Hyg. 1995;52:565–8.

19. Hofmann N, Mwingira F, Shekalaghe S, Robinson LJ, Mueller I, Felger I. Ultra-sensitive detection of Plasmodium falciparum by amplification of multi-copy subtelomeric targets. PLoS Med. 2015;12:e1001788.

20. Li H. Aligning sequence reads, clone sequences and assembly contigs with BWA-MEM. arXiv preprint 1303 3997 [Internet]. 2013; Available from: http://arxiv.org/abs/1303.3997

21. Li H, Handsaker B, Wysoker A, Fennell T, Ruan J, Homer N, et al. The Sequence Alignment/Map format and SAMtools. Bioinformatics. 2009;25:2078–9.

22. Auwera GAV der, Van der Auwera GA, Carneiro MO, Hartl C, Poplin R, del Angel G, et al. From FastQ Data to High-Confidence Variant Calls: The Genome Analysis Toolkit Best Practices Pipeline [Internet]. Current Protocols in Bioinformatics. 2013. p. 11.10.1–11.10.33. Available from: http://dx.doi.org/10.1002/0471250953.bi1110s43

