## Supplemental_Table_1_and_2 for "Optimization of whole-genome sequencing of *Plasmodium falciparum* from low-density dried blood spot samples"

**Supplementary tables**

**Table S1. PCR conditions for sWGA amplification using modified sWGA primers.**

| **Reagents** | **Volume**  **(for 50μL reaction)** | **Cycling condition** |
| --- | --- | --- |
| 10X NEB Phi29 Buffer | 5 μL | 35°C for 5 min  34°C for 10 min  33°C for 15 min  32°C for 20 min  31°C for 30 min  30°C for 16 hrs  65°C for 15 min  10°C hold |
| 10 mg/ml BSA | 0.5 μL |  |
| 100μM Primer set 6A* | 1.25 μL |  |
| 250μM Primer set 10A* | 0.5 μL |  |
| 5mM dNTP (Adjusted 70%AT: 30%GC) | 19.75 μL |  |
| NEB Phi29 Enzyme | 3 μL |  |
| Template DNA | 20μL |  |
| **Total** | **50μL** |  |

*Primer sequences were previously reported for 6A (Sundararaman *et al.,* 2016) and 10A (Oyola *et al.,* 2016).

**Table S2. Estimates of parasite densities from extracted DBS and WGA reactions**

| **Parasites per μL of blood** | **Parasitaemia*** | **# of parasites per punch**** | **# parasite per μL of extracted DNA** | **# parasites per WGA reaction** |
| --- | --- | --- | --- | --- |
| 10 | 0.0002 | 120 | 0.8 | 16 |
| 100 | 0.002 | 1200 | 8 | 160 |
| 1,000 | 0.02 | 12000 | 80 | 1600 |
| 10,000 | 0.2 | 120000 | 800 | 16000 |

*Parasitaemia = 100X(number of parasites per μL of blood/5 million RBC per μL of blood)

**An estimated average size of DBS is ~10mm in diameter, a 6mm punch contains ~60% of the 20μL of blood spotted on the Whatman paper
